# Genome-wide Genetic Marker Analysis and Genotyping of *Escherichia fergusonii* strain OTSVEF–60

**DOI:** 10.1101/2020.07.19.209635

**Authors:** Otun Saha, Nadira Naznin Rakhi, M. Nazmul Hoque, Munawar Sultana, M. Anwar Hossain

**Affiliations:** Department of Microbiology, University of Dhaka, Dhaka 1000, Bangladesh; Department of Biotechnology and Genetic Engineering, Bangabandhu Sheikh Mujibur Rahman Science and Technology University, Gopalganj, Bangladesh; Department of Gynecology, Obstetrics and Reproductive Health, Bangabandhu Sheikh Mujibur Rahman Agricultural University, Gazipur-1706, Bangladesh

**Keywords:** Whole genome sequence, *Escherichia fergusonii*, poultry, virulence, multidrug, metal resistance

## Abstract

Poultry originated *Escherichia fergusonii* (POEF), an emerging bacterial pathogen, causes a wide range of intestinal and extra-intestinal infections in the poultry industry which incurred significant economic losses worldwide. Chromosomal co-existence of antibiotics and metal resistance genes has recently been the focal point of POEF isolates besides its pathogenic potentials. This study reports the complete genome analysis of POEF strain OTSVEF-60 from the poultry originated samples of Bangladesh. The assembled draft genome of the strain was 4.2 Mbp containing 4,503 coding sequences, 120 RNA (rRNA = 34, tRNA = 79, ncRNA = 7), and three intact phage signature regions. Forty one broad range antibiotic resistance genes (ARGs) including *dfr*A12, *qnr*S1, *bla*_TEM-1_, *aad*A2, *tet*(A) and *sul*-2 along with multiple efflux pump genes were detected, which translated to phenotypic resistant patterns of the pathogen to trimethoprim, fluoroquinolones, β-lactams, aminoglycoside, tetracycline, and sulfonamides. Moreover, 22 metal resistance genes were found co-existing within the genome of the POEF strain, and numerous virulence genes (VGs) coding for *cit* (AB), *feo* (AB), *fep* (ABCG), *csg* (ABCDEFG), *fli*C, *omp*A *gad*A, *ecp*D etc were also identified throughout the genome. In addition, we detected a Class I integron gene cassette harboring *dfr*A12, *ant* (3″)-I and *qac*EΔ-*sul*2) genes, 42 copies of insertion sequence (IS) elements, and two CRISPR arrays. The genomic functional analysis revealed overexpression of several metabolic pathways related to motility, flagellar assembly, epithelial cell invasion, quorum sensing, biofilm formation, and biosynthesis of vitamin, co-factors, and secondary metabolites. We herein for the first time detected multiple ARGs, VGs, mobile genetic elements, and some metabolic functional genes in the complete genome of POEF strain OTSVEF-60, which might be associated with the pathogenesis, spreading of ARGs and VGs, and subsequent treatment failure against this emerging avian pathogen with currently available antimicrobials.

## Introduction

*Escherichia fergusonii*, an opportunistic emerging pathogen is the closest relative of *E. coli* showing 64% similarity in DNA hybridization^1^. This species was formerly known as Enteric Group 10, due to its biochemically distinct nature compared to other species, and bio-groups of the family *Enterobacteriaceae*^2^. *E. fergusonii* was initially isolated from human clinical samples^2,3^, however can cause septicemia and diarrhea in animals including several avian species^4-7^. The first available whole genome data of *E. fergusonii* was published as a part of survey analyzing *E. coli* genome evolution. Since the first report, there has been an increasing number of case reports of *E. fergusonii* of animal origin, isolated mainly from feces of domestic animals and birds^4-7^. This organism is also being frequently isolated from foods of animal origin such as beef, and cheese during routine screening^9^. But its significance regarding public health concern was not realized initially, while it has recently been considered as an emerging zoonotic pathogen^2,3^, and also possesses an extended spectrum of resistance to antibiotics^10,11^. This enteric pathogen can cause a variety of clinical symptoms in poultry including septicemia and diarrhea^4,7^ with an approximate mortality rate of 18–30% in one-day-old chicks^11^. This bacterium was also identified as the dominant member of the aerobic microbial community in birds including poultry^4,12^.

The emergence of antimicrobial resistant, and pathogenic strains of bacteria in food-producing animals especially the chicken represents a serious food safety concern, and is a threat to public health. The determinants of antibiotic resistance have been identified on mobile genetic elements (MGEs), such as plasmids, transposons, and integrons of the bacteria isolated, identified and sequenced from poultry originated samples, thus potentially allowing these determinants to be spread between species^11,13^. The plasmids of bacteria are self-replicating, extrachromosomal replicons, and crucial agents of genetic change in microbial populations. In addition to conferring resistance to multiple antibiotics, naturally occurring plasmids can also promote the spread of a variety of traits, including resistance to mercury and other heavy metals, virulence, fitness, and the metabolism of unusual compounds^13,14^. The extended spectrum of antibiotic resistance genes (ARGs), and its association with MGEs in *E. fergusonii* isolates has mainly attracted attention to this bacterial species in recent times^11^. In a recent study, Simmons et al. (2016) reported that the majority of the isolated POEF were resistant to β-lactams, aminoglycosides, tetracycline and sulfonamides in different broiler chicken farms of Canada^15^. Besides, several other studies on POEF reported plasmid associated resistance markers responsible for resistance to aminoglycosides, tetracycline, sulfonamide, trimethoprim and beta-lactams as well as the chromosomal mutation on *gyr*A mediated quinolone resistance^16,17^. Therefore, the association of bacterial MGEs with antimicrobial resistance might have contributed to the dissemination of ARGs and VGs among multiple bacterial genera of human, and veterinary importance^14^. Furthermore, the presence of ARGs carried by MGEs, such as plasmids, may also lead the resistance genes to enter into the human food chain through horizontal gene transfer to other enteric bacteria or bacteria of potential food safety concern^18^. However, operons related to tolerance to heavy metals including manganese, mercury, nickel, tellurium, silver, copper and cobalt-zinc-cadmium were also found in *E. fergusonii* of poultry origin^16^. Most importantly, the mobilome carrying heavy metal tolerance operons may lead to the co-selection of antibiotic-resistance, when heavy metals in animal feeds (copper, zinc and nickel for improvement of animal health and growth), or in topical medications (silver and copper for the treatments of superficial wounds) or the feed contaminants (mercury and arsenic) reach a toxic concentration causing a long term selective pressure^19^.

Therefore, the pathogenic potentiality along with resistance determinants makes this organism a potential threat not only to animal health, but also to public health. Till to date, only few pathogens such as *Salmonella, Enterococcus* spp., *Clostridium perfringens* and *Escherichia coli* from poultry have been studied extensively at genomic level^16^. In order to investigate the role of *E. fergusonnii* in the pathophysiology of poultry as well as its contributions in spreading ARGs the reported superbug *E. fergusonii* strain OTSVEF-60 was isolated from poultry farm of Bangladesh. Here we report the whole genome of the superbug and functional genomics in order to understand the mechanisms how this superbug showed its resistant phenotypes, contributes in ARGs spreading and pathogenesis.

## Materials and Methods

### Bacterial isolate

POEF OTSVEF-60 strain was isolated from a dropping sample collected from Savar District of Bangladesh (23.85° N, 90.26° E). Followed by retrieval of the isolate on Nutrient agar plate, the isolate was subjected to Gram staining and different biochemical tests such as catalase test, oxidase test, fermentation of glucose, lactose and sucrose, urease, oxidase, catalase, methyl red, Voges-proskauer, indol test, Simon’s citrate and hydrogen sulfide according to the guideline of the Bergey’s manual of Determinate Bacteriology^20^.

### Biofilms formation (BF) assay

The BF assay of POEF-OTSVEF-60 strain was performed using 96 well microtiter plate method for quantitative analysis of attachment, and BF on plastic surfaces in static conditions^21^. The assay was performed in duplicate in the 96 well tissue culture plates. The observed optical density (OD) was evaluated to determine the biofilm-forming ability of the isolates on a 4-grade scale (non-adherent, weakly adherent, moderately adherent, and strongly adherent). These four OD grades were determined by comparing OD with ODc (three standard deviation values above the mean OD of the negative control). Furthermore, the biofilm surface was stained by Film tracer LIVE/DEAD biofilm viability kit after 24 hours to observe the proportion of live or active cells (fluorescent green), and dead or inactive cells (fluorescent red) using fluorescence microscope and DP73 digital camera (40X objective).

### Antimicrobial susceptibility testing

The isolate was subjected to antibiotic susceptibility testing by the Kirby-Bauer agar disc diffusion method to determine their multidrug (MDR) pattern. The tested antibiotics included 14 antimicrobial drugs belonged to 12 antibiotic groups: Penicillins (ampicillin, Amp-10 μg); Tetracyclines (doxycycline, Do-30 μg; tetracycline, Te-30μg); Nitrofurans-(nitrofurantoin, F-300 μg); Lipopeptides (polymexin B, Pb-30 μg); Quinolones (subclass fluoroquinolone ciprofloxacin, Cip-10 μg; subclass quinolone, nalidixic acid, Na-30μg); sulfonamide (sulfonamide, Sul-250μg); trimethoprim, TM-5μg; Penems-(imipenem, Imp-10 μg); Aminoglycosides (gentamycin, GN-10 μg); Phenols (chloramphenicol, C-30 μg); Cephems (oral)-(cephalexin, Cex-30 μg); Macrolides (azithromycin, Atm-15 μg), which are widely used in poultry farms^22^.

### Genomic DNA extraction, whole genome sequencing and genome assembly

To extract the genomic content of the POEF strain OTSVEF-60, a commercial DNA extraction kit, QIAamp DNA Mini Kit (Qiagen, Hilden, Germany) was used according to the manufacturer’s instruction. The harvested DNA was visualized on 1% (w/v) agarose gel, and DNA concentration and purity was measured by a NanoDrop 2000 UV-Vis Spectrophotometer (Thermo Fisher, Waltham, MA, USA). Whole genome sequencing was performed under Ion-Torrent High Throughput sequencing platform using SeqStudio Genetic Analyzer^23^.

Machine generated data was transferred to the Ion Torrent server, where data were processed through signal processing, base calling algorithms, and adapter trimming to produce mate-pair reads in FASTQ format. The FASTQ reads quality was assessed by the FastQC tool^24^ followed by trimming of low quality reads and reads less than 200 bp using the Trimmomatic tool^24^, while the quality cut off value was Phred-20. Reads were assembled with the SPAdes, (version 3.5.0) genome assembler^25^. Generated assembled reads were mapped to, and reordered according to a reference sequence of *E. fergusonii* ATCC 35469 complete genome from NCBI (accession number: CU928158) by progressive Mauve algorithm in Mauve software^26^.

### Sequence analysis

Draft genome of POEF OTSVEF-60 strain was initially annotated using RAST (Rapid Annotation using Subsystem Technology) server^27^, and KEGG (Kyoto Encyclopedia of Genes and Genomes) Automatic Annotation Server (KAAS)^28^. The RAST server provided data on the distribution of genes in various categories. Besides, the circular visualization and comparison showing the distinct differences between multiple genomes were performed by BLAST Ring Image Generator (BRIG)^29^ with set default parameters. The reference genomes for comparison were retrieved from the GenBank database (http://www.ncbi.nlm.nih.gov/genbank/). To determine the average nucleotide identity (ANI), the comparison between the available genomes were performed using JSpeciesws database (http://jspecies.ribohost.com/jspeciesws) which used Blast alignments to evaluate whole genome homologies. Cut-off values for species delineation were set to 95% ANI on 69% of conserved DNA^30^. We employed the tRNAscan^31^ and rRNAmmerv1.2^32^ tools for the detection of tRNA and rRNAs in the genome. Besides, all contigs were submitted to the CGE Finder Series (Centre for Genomic Epidemiology, Technical University of Denmark (DTU) (https://cge.cbs.dtu.dk/services/) to extract the species, multilocus sequence type (ST), pathogenic level, plasmid replicon types along with information about genes mediating resistance to β-lactams and fluoroquinolones, and distinct virulence genes using the tools SpeciesFinder 1.3, MLST 1.8, PathogenFinder 1.1, PlasmidFinder 1.3, ResFinder 2.1, VirulenceFinder 1.5), respectively^33^. The total number of prophage sequences was determined using Phage search tool enhanced release (PHASTER; http://phaster.ca/)^33^. The tool identifies intact, questionable, and incomplete prophage sequences by scores of >90, 70–90, < 70, respectively. Moreover, clusters of regularly interspaced short palindromic repeats (CRISPR)-Cas system of POEF OTSVEF-60 strain was characterized based on annotation by CRISPR one, a web-based tool (http://omics.informatics.indiana.edu/CRISPRone), which provides class, type, and subtype of CRISPR-Cas system, and number, length and nucleotide sequences of repeats and spacers^34^. The insertion sequence (IS) elements were predicted using the IS Finder (https://www-is.biotoul.fr)^35^.

To reveal the presence of antimicrobial resistance markers (AMRs) in POEF OTSVEF-60, we mapped the genomic data against CARD 3.0.0 (The Comprehensive Antibiotic Resistance Database) database, a rigorously curated collection of known resistance determinants and associated antibiotics. Furthermore, secondary metabolite synthesizing genes were identified by genome mining pipeline of AntiSMASH 5.0^36^. However, NCBI Tree Viewer (https://www.ncbi.nlm.nih.gov/tools/treeviewer/) and MEGA v7.0^37^ was used for performing whole genome based phylogenetic analysis applying Neighbor-Joining method with 1,000 bootstraps. The phylogenetic tree was subsequently imported to iTOL (v. 3.5.4) (http://itol.embl.de/)^38^ for better visualization, and bootstrap values were reported for each branch.

### Nucleotide sequence accession numbers

The complete genome sequence, annotated POEF OTSVEF–60 chromosome, and integron sequences have been deposited in NCBI GenBank under the accession numberJABCUC000000000.

## Results

The whole genome sequence (WGS) analysis of the study isolate, OTSVEF–60 using Kmer Finder 3.1 identified the isolate as *E. fergusonii*, while the pathogenicity of the isolate was confirmed (0.936 out of 1.00 (near the pick value indicating higher pathogenicity)) by PathogenFinder 1.1, as also supported by the biofilm assay (OD=0.431). Besides, the isolate was found as multidrug resistant (MDR) showing resistance to antibiotic: ampicillin, sulfonamide, trimethoprim tetracycline, doxycycline, gentamycin, ciprofloxacin, nalidixic acid according to the antibiotic susceptibility assay.

### Genome assembly and annotation

The WGS of POEF strain OTSVEF–60 revealed a circular chromosome spanning 4,661,770 bp with an average GC content of 49.8%. This WGS possessed 263 contigs with two incomplete plasmids (IncX1 and p0111) in the genome (Table 1). According to RASTtk annotation, only 37% of the genes were located in the generated subsystems, and majority (63%) of the annotated genes remained out of the subsystem list. The genome was found to bearing 4,503 coding sequences (CDS) according to RAST annotations. Annotated results from subsystems, and pathway reconstruction are depicted in Fig. 1.

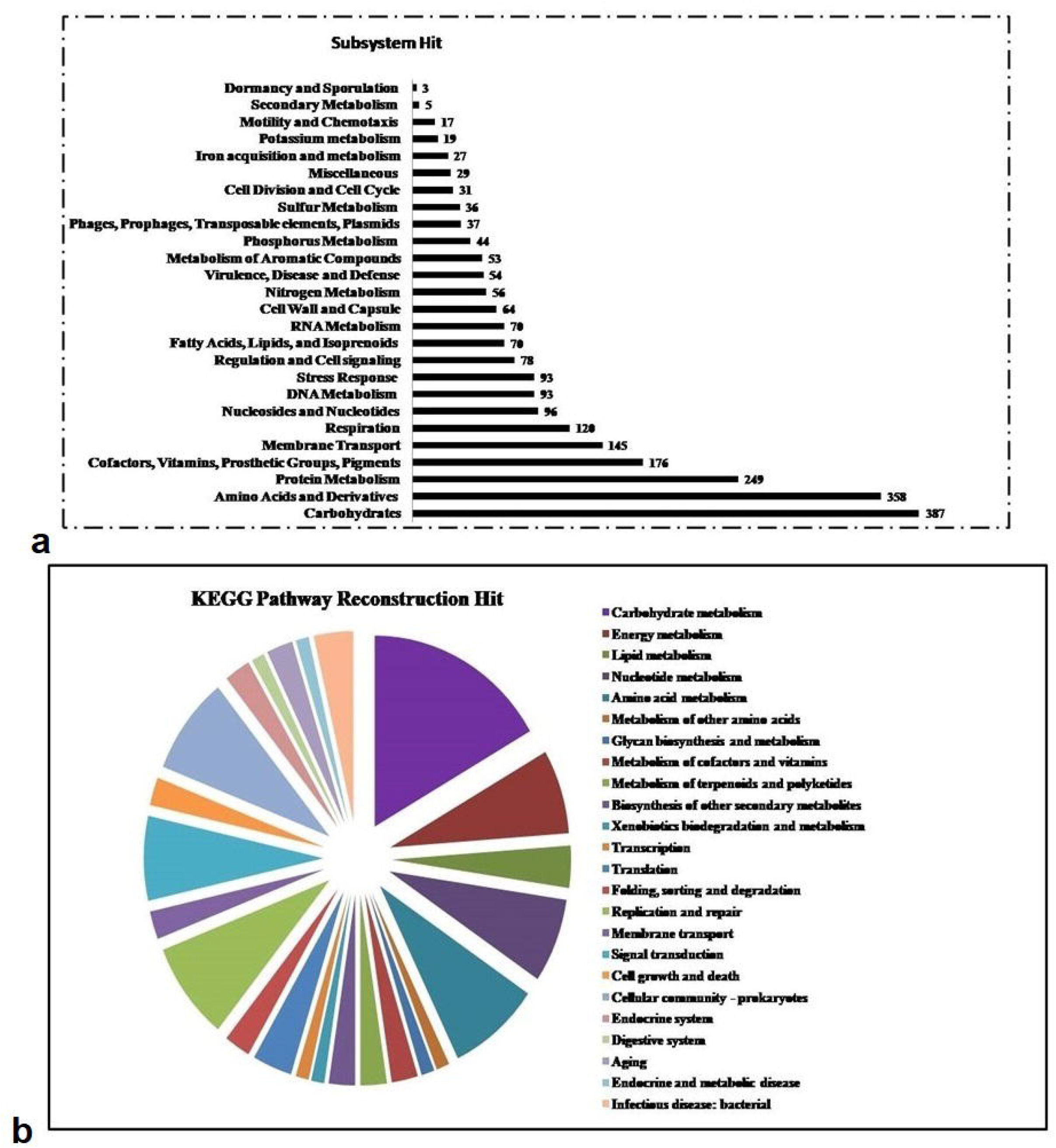

Although, the BLAST and SEED close strain analyses revealed that the genome had similarity with POEF along with other *Escherichia coli* strains, the Kmer-based genomic comparison, and whole genome-based phylogeny showed that the POEF strain OTSVEF–60 had the closest proximity to the strain *E. fergusonii* ATCC 35469 (Fig. 2). All the *E. fergusonii* (CU928158.2-CP042942.1) strains carrying the same flagellar type were clustered together (Fig. 2). Additionally, *E. coli* strain (CP027579.1-CP040919-1) was distantly related to other serotypes, and clustered separately (Fig. 2). *In silico* analysis of the genome using housekeeping genes (*adk, fum*C, *gyr*B, *icd, pur*A and *rec*A) which clustered with the human strains revealed that this strain might belong to a new ST-5637. The genetic features of POEF strain OTSVEF-60 complete genome assembly, and annotations are summarized in Table 1, and circular graphical map of the genome is shown in Fig. 3 and Fig. 4. We did not find any similarities to the allelic profile of any other *Escherichia* strains including *E. fergusonii* for *fum*C and *icd* genes (Fig. 3, Supplementary Table 1). The screening for acquired antibiotic resistance genes revealed the presence of different resistance genes in the chromosome of POEF OTSVEF-60 isolate (Fig. 4, Table 1). Classification, general features, and genome sequencing information of POEF-OTSVEF-60 according to the MIxS recommendations are presented in Table 2.

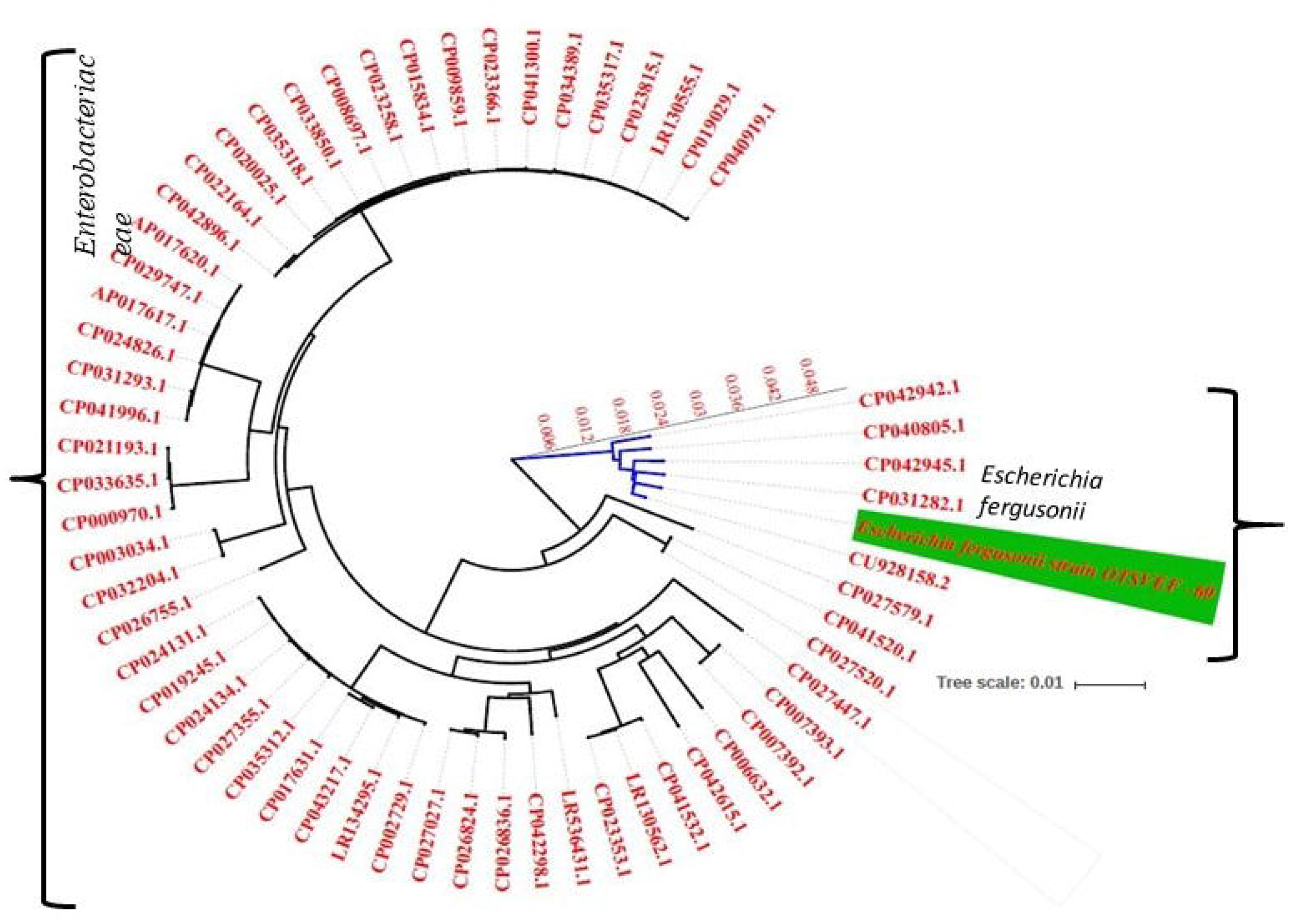

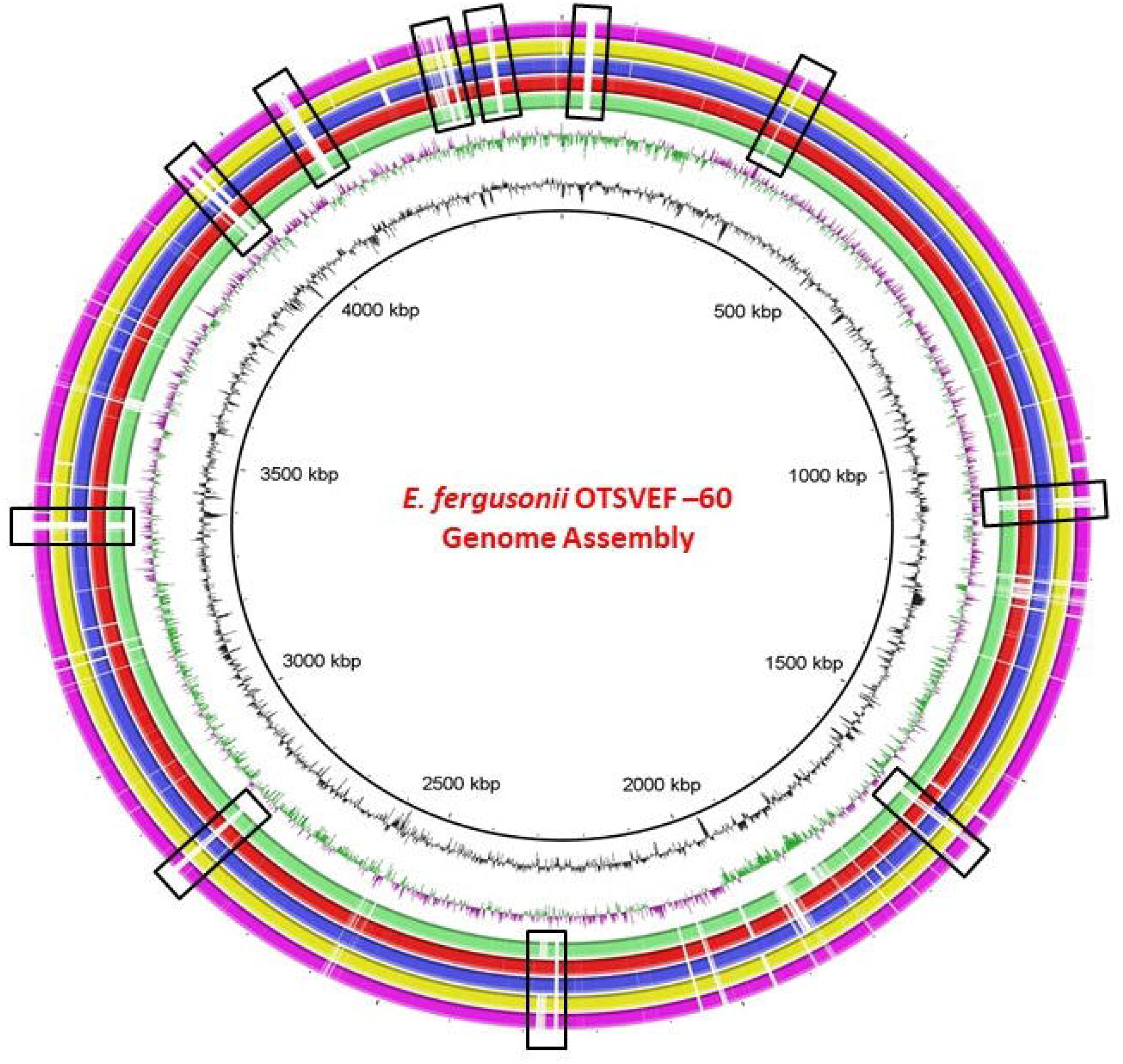

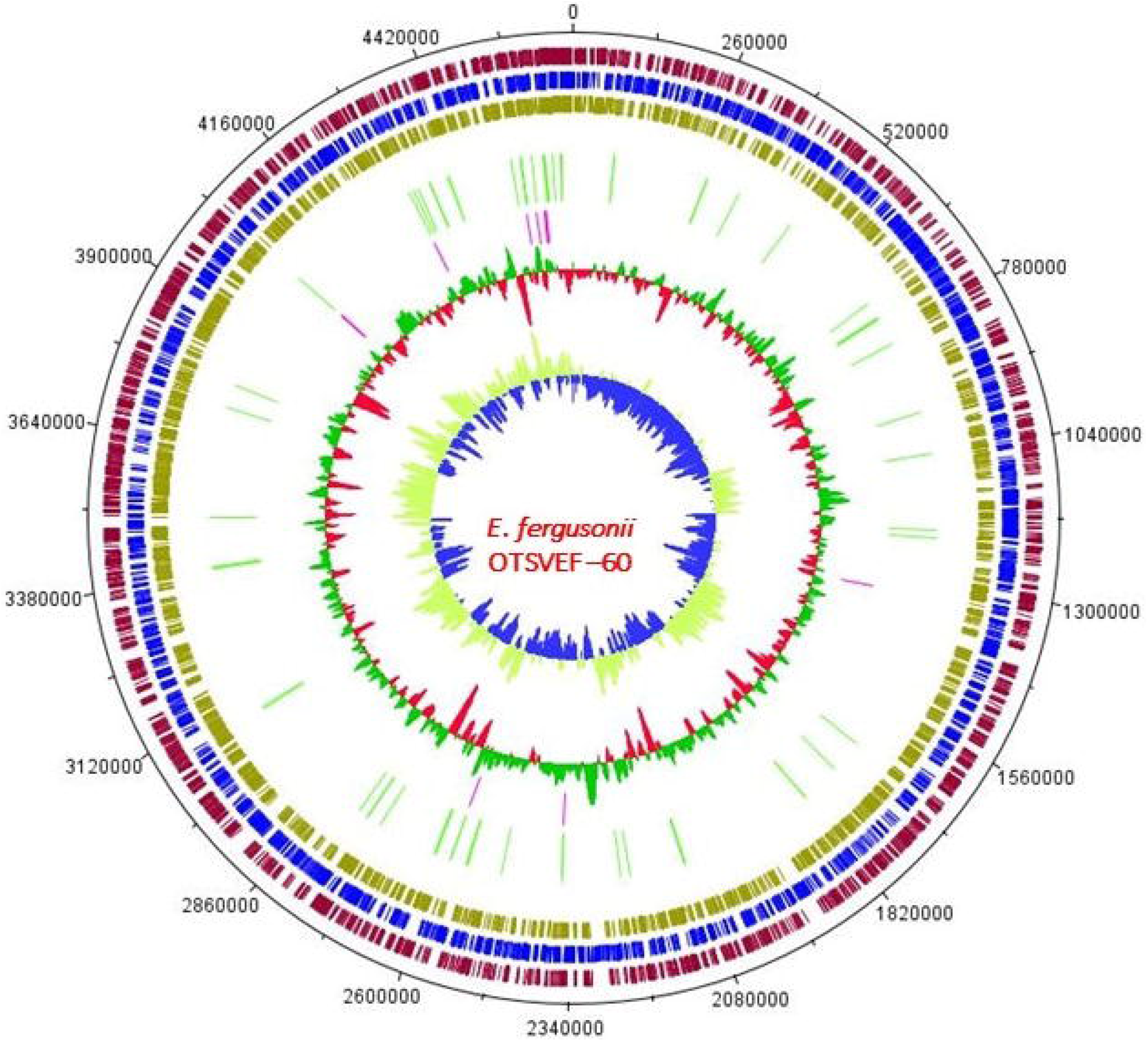

### Identification of prophages in POEF-OTSVEF-60 genome

We identified six prophages in the WGS of the POEF-OTSVEF-60 strain, of which lambda-like prophages were predominantly abundant. The OTSVEF-60 strain carried three intact prophage sequences (region 1, 2, 5) along with two incomplete (region 4 and 6), and one questionable prophage (region 3) sequences (Fig. 5). The resemblance of the intact prophage sequences was found with Salmon_ sal5 (region 5), and Entero _ fiAA91 (region 1), and Entero_mEp460 (region 2). Besides, the incomplete prophage region 4 resembled the Entero_P4 (Fig. 6, Table 3). All of the prophage sequences gave significant (coverage > 80%, identity > 90%) hits against NCBI-GenBank reference sequences. Most of the deduced amino acid sequences of phages best matched either with phage proteins or with hypothetical proteins (Table 3).

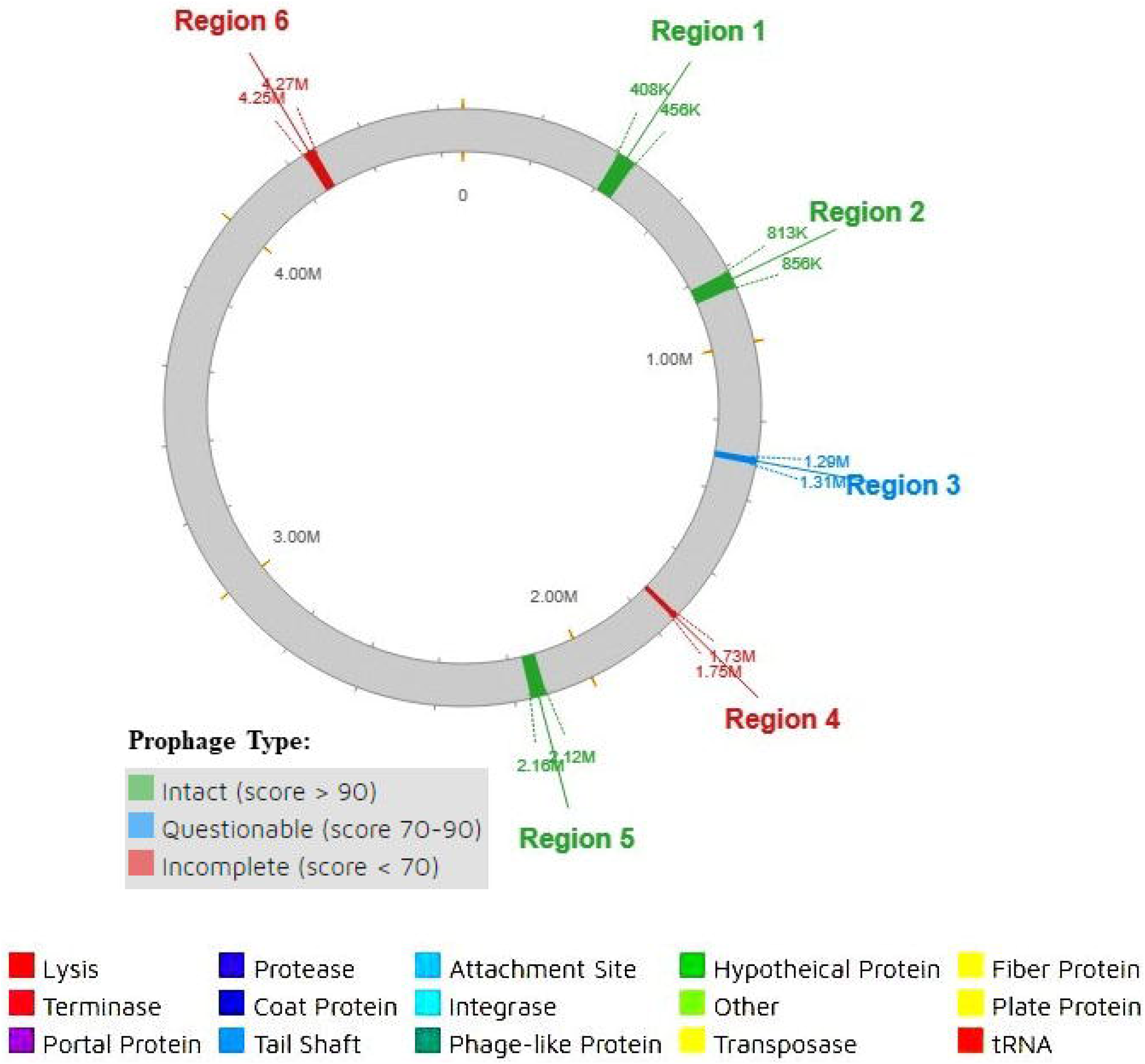

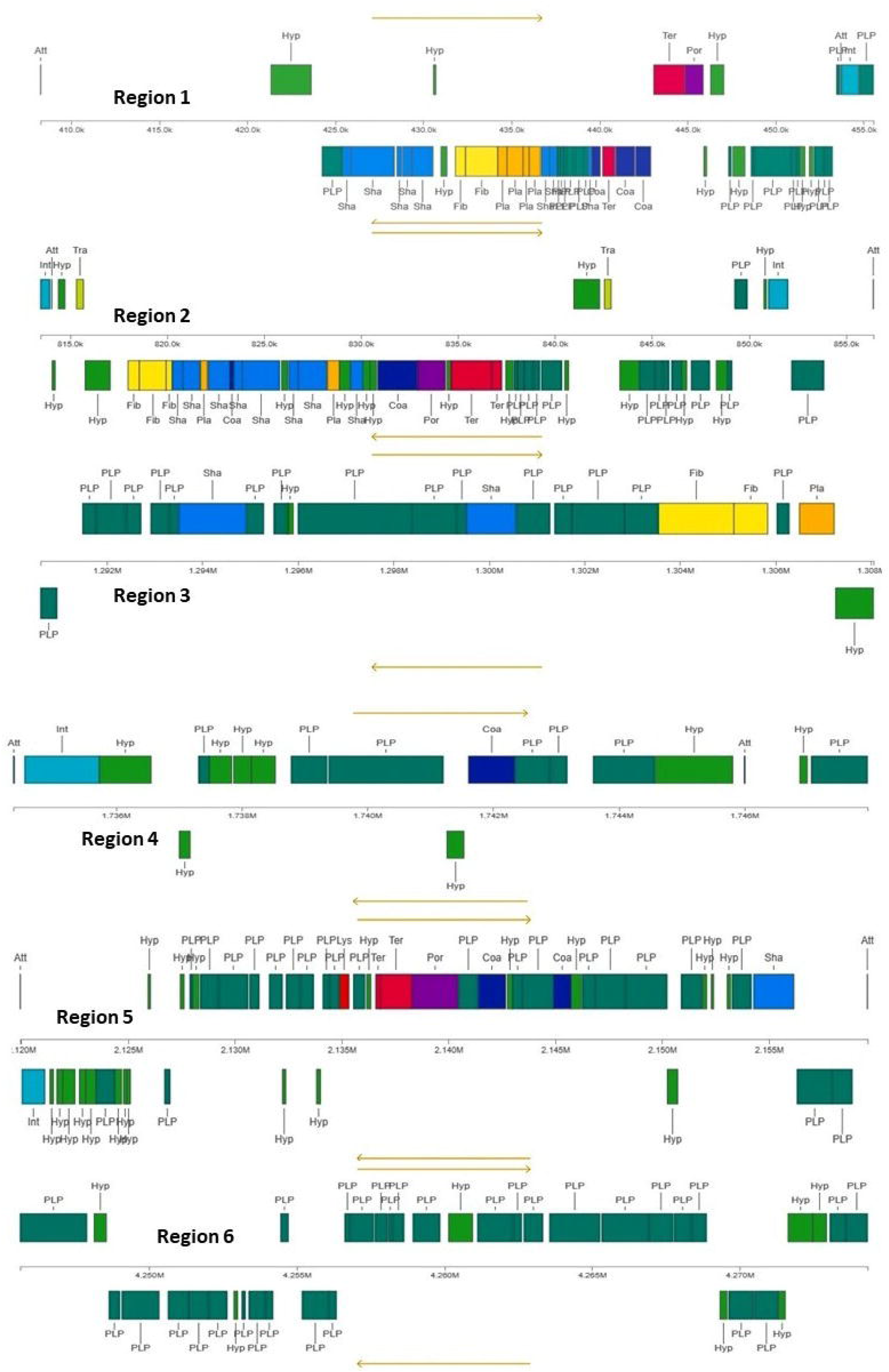

### CRISPR-*Cas* system of POEF-OTSVEF-60 strain

The most common subtype of CRISPR-*Cas* system found in the POEF-OTSVEF-60 was I-A, and other subtypes identified were type-1-E. Thus, the isolate carried two separate CRISPR arrays which were segregated into 3 CRISPR regions. Among those 3 CRISPR regions, one of the CRISPR region lacked *cas* genes whereas remaining two CRISPR regions carried three or more *Cas* encoding genes. However, repeats and spacer units were not conserved into the POEF-OTSVEF-60 genome (Supplementary Table 2).

### Antimicrobial resistance analysis

There was a wide distribution of antibiotic resistance genes (ARGs) including both antibiotic inactivating enzyme and efflux pump encoding genes in the genome of POEF OTSVEF-60. These ARGs comprised 41 genes conferring resistance to a wide range of antimicrobials including trimethoprim, fluoroquinolones, β-lactams, aminoglycoside, tetracycline, and sulfonamides. RAST analysis identified 13 AMR gene markers including *aadA2, kdp*E (resistance to aminoglycoside); the non-ESBL gene *bla*_TEM-1_ (penicillin); *dfr*A12 (trimethoprim); *qnr*S1, *par*C and *gyr*B (resistance to fluoroquinolone); *sul1, sul2* (sulfonamide); and *mph*A (macrolide) conferring resistance to different groups of antibiotics which were also consistent with the findings of in vitro antibiotic susceptibility tests (Supplementary Fig. 1, Table 6). Besides, efflux pump genes such as *tet*A *emr*A, and *emr*B conferring resistance to tetracyclines, and fluoroquinolone, respectively were also found. Several other efflux systems such as multidrug facilitator superfamily, resistance-nodulation-division superfamily efflux, and gene corresponding to the ATP-binding cassette superfamily were also detected in this complete genome.

The *in vitro* antimicrobial resistance profiling also suggested the possible expression of the ARGs detected in in silico analysis. (Table 6). Besides, chromosomal point mutations (TCG ⍰ TTG/S ⍰ L) in the *gyr*A gene might be responsible for resistance to 4^th^ generation ciprofloxacin, and nalidixic acid. However, the isolate showed susceptibility to some of the tested antimicrobials including polymyxinB, nitrofuran, and chloramphenicol. Furthermore, a classical class 1 integron containing a *qacEΔ-sul1* element for disinfectant, and sulfonamide resistance was retrieved within the contig 57 of the complete genome. The variable region of this integron also harbors ant (3″)-I, and *dfrA* gene cassettes (Fig. 7a). On the other hand, the beta-lactam resistance gene, *bla*_TEM-1_ was flanked by *qnr*B gene encoding *Qnr*B protein, which is a member of the pentapeptide repeat protein (PRP) family, and has been shown to block the action of ciprofloxacin on purified DNA gyrase and topoisomerase IV (Fig. 7b).

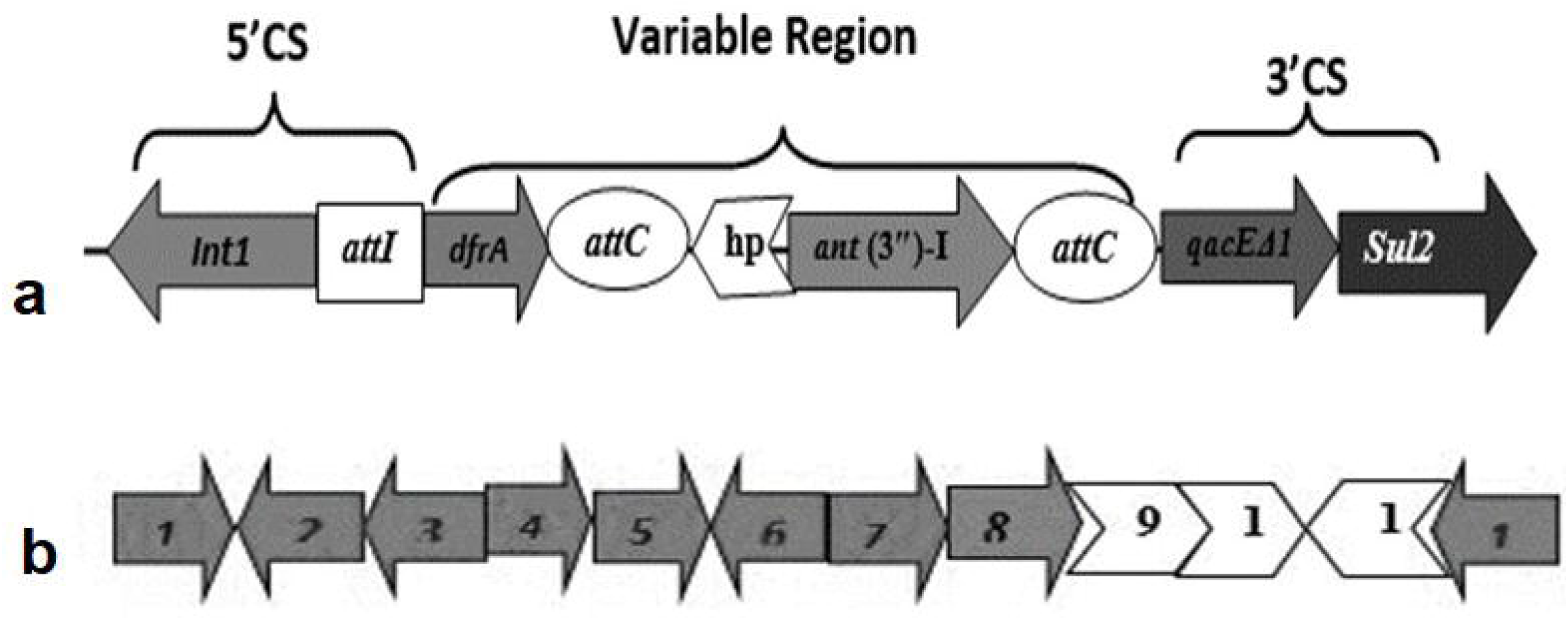

### Metal resistance genes across the genome of POEF-OTSVEF-60

We simultaneously identified 22 heavy metal resistance genes (HMRGs) across the genome of POEF-OTSVEF-60 including copper (*cue*R), zinc (*Zra*P), chromium (*chr*A), and arsenate (*ars*R, *ars*B, *ars*C) genes. In addition to copper resistance gene, both copper homeostasis related genes (*cue*O, *cop*D, *cop*C, *cup*C-D), and copper tolerance genes (*cut*A,*cut*E,*cut*F,*cut*C) were also present in this genome. Moreover, the isolate harbored seven chromosome-encoded cobalt-zinc-cadmium resistance genes namely *zit*B, *cus*A, *cus*F, *cus*C, *czc*D, *cus*R *cus*S, and genes coding for ferrous-iron efflux pump (*fie*F), nickel responsive regulator (*nik*R), and molybdenum transport system binding proteins (*mod*A, *mod*B, *mod*C, *mod*F) (Table 4).

### Virulence analysis

VGs were determined in silico using reference literature, BLAST and Virulence finder at 96% identity. Virulence genes (VG) with more than 95% identity were considered positive in this genome analysis. In the genome of POEF-OTSVEF-60, 39 VGs were identified, including those encoding citrate two-component system and lyase, *cit*(AB); ferrous iron transport, *feo*(AB); ferrienterobactin uptake, ABCDGE;, Tol system, *tol* (ABCQR); nucleoside-specific channel-forming,*tsx*; curli production component, *csg* (ABCDEFG); flagellin, *fliC*;, glutamate decarboxylase, *gad;*, outer membrane porin, *omp*A;, arginine ABC transporter, *art*J; enterobactin synthetase component, *ent*(CEFS); fimbria adhesion, *csp*D and two putative virulence factors (*mvi*M, *vir*K) (Table 5). This profile shared similarities with the closely related strain *E. fergusonii* ATCC 34569 except *tol*A, *csg* (ABCDFG) (Table 5). We found no plasmid-encoded VGs in the study genome.

### Metabolic functional potentials of the genome

Metabolic pathway analysis of POEF-OTSVEF–60 genome revealed a number of complete and incomplete pathways possibly associated with the AMR, and virulence properties of the isolate (Fig. 1). The detected functional pathways were distributed under five major KEGG orthologies (KO) such as cellular process, metabolism, environmental information processing, genetic information processing, and human diseases (Fig. 1). A deeper look into these KEGG pathways disclosed that they were associated with the motility, flagellar assembly, quorum sensing (*bap* A, *lsr*K), biofilm formation (*cya*A, *glg*P, *acf*D); biosynthesis of vitamins and co-factors, folate (*fol1*), and one carbon pool (*met*H, MTR). Moreover, genes coding for secondary metabolites (*rfb*A, *rff*H) such as streptomycin, acarbose, and validamycin, xenobiotic degradation (*pob*A) related to amino benzoate degradation were also identified in this genome. We also detected several gees (*nap*A, *pob*A, *nar*G, *nar*Z, *nxr*A, *nir, glt*B, *frd*A, *suc*A) that are associated with microbial metabolism, and adaptation of the isolate in diverse environments. The gene (*yee*J) responsible for the bacterial invasion of epithelial cells leading to the adhesion to different abiotic surfaces, and promoting biofilm formation was also identified. Furthermore, the AntiSMASH tool detected two secondary metabolite gene clusters in the study genome, which encoded arylpolyene (protective agents against oxidative stress), and non-ribosomal peptide synthase (NRPS) (toxins, siderophores, pigments, antibiotics, cytostatics, immunosuppressant’s or anticancer agents) that support the bacteria for surviving in harsh environment (Supplementary Table 3, Supplementary Fig. 2).

### Insertion sequence (IS) elements

Forty-two copies of IS elements were identified in the OTSVEF–60 genome (Supplementary Table 2). The IS Finder software identified 10 families of IS elements, of which IS3 and Tn3 were most predominant followed by ISKra4, IS200/IS605, IS110, ISAs1, IS5, IS6, IS1, IS4 family. Moreover, 23 truncated transposases were also found in the OTSVEF-60 genome (Supplementary Table 4).

## Discussion

This study to the best of our knowledge, reports the first isolation and characterization multidrug resistant *E. fergusonii* of clinical importance from the poultry samples of Bangladesh. POEF is the emerging zoonotic pathogen that have been reported in both humans and animals globally^3,4,11,12^. Probably the species has been mistaken as *E. coli* until now, as typical 16S rRNA gene sequencing based identification method frequently failed to distinguish it from *E. coli*^39^. The growing concern of this organism as a zoonotic pathogen insisted to analyze the whole genome of the poultry isolate superbug *E. fergusonii* OTSVEF-60 for understanding its pathogenicity and antimicrobial resistance.

POEF-OSSVEF-60 possessed more VGs compared to the non-pathogenic *E. fergusonii* strain ATCC 35469. These VGs comprised of both chromosome and MGEs including plasmids, prophage-like elements, and genomic islands. The VGs also included various curli fimbriae genes to be associated with virulence, and biofilm formation (BF)^40^. Moreover, the Flagellin, adhesion, fimbria adhesion, and BF regulatory genes were detected in the genome of POEF-OSSVEF-60 which might also contribute to the virulence of the strain, and subsequent treatment failure leading to antimicrobial resistance to a variety of drugs (being MDR)^10^. Moreover, these virulence properties shared similarities with the closely related strain *E. fergusonii* ATCC 34569 except *tol*A, *csg* (ABCDFG)^8,11^. Moreover, the observed combinations of VGs might be involved in a distinct category of pathogenic *E. fergusonii*, which signifies its possible role behind the outbreak in the poultry farm.

The results of the *in vitro* resistance assay corroborated with the genomic resistance analysis showing the strains as a MDR (resistant to penicillin, sulfonamide, tetracycline, aminoglycoside, and fluoroquinolone)-pathogen. In this study, *bla*_TEM-1_, *aad*A2, *kdp*E *qnr*S1 were the most dominant ARGs type which was consistent with other studies^41,42^. Besides the ARGs of the study isolate also included efflux pumps mediated resistance genes: resistance nodulation division (RND) type, ATP binding cassette (ABC), and major facilitator superfamily (MFS) of multidrug transporters. The genes of RND type efflux pump proteins were up-regulated, while *mdt*ABC-mediated novobiocin resistance^43^ along with decreased susceptibility to erythromycin, fosfomycin and fluoroquinolone^44,45^ have also been reported. The tetracycline-resistant gene, *tet*A deserves particular attention as the tetracycline resistance genes are evident of being physically linked to other resistance genes^15^. However, the isolate was susceptible to polymyxinB, nitrofuran, and chloramphenicol, these finding are in line with the findings of Simmons et al. who also reported that *E. fergusonii* was susceptible to imipenem, and resistant to ciprofloxacin^15^. In another study, Forgetta et al. reported that an *E. fergusonii* strain was resistant to 22 antibiotics supporting our current findings^11^. Interestingly, unlike the previous and first report of the *E. fergusonii* whole genome, along with antibiotic resistance genes, multiple metal resistance and tolerance genes conferring resistance to copper, chromium, iron, potassium, cobalt, nickel, and zinc were found in the OTSVEF–60 strain. Metals have been found to interfere with several bacterial cellular functions such as protein activity, oxidation, nutrient assimilation, membrane stability, DNA replication^21,46^ along with contributing to the cross-resistance to antibiotics^47^. This co-existence might be due to the selective pressures caused by heavy metals in the environment which indirectly induces the selection of antibiotic resistance, especially in the environment contaminated with these two elements^47^ or vice-versa. Moreover, the metal resistance genes can also increase the AMR and virulence of the bacterial pathogens^48,49^. The study strain also carried numerous MGEs including class I clinical origin integronI (*int*I) gene cassette with resistance genes for aminoglycoside, sulphonoamide and trimethoprim. Presence of *int*I gene cassette makes this superbug capable of disseminating antibiotic resistance to other bacteria^50^. Yet to date, very limited literature is available regarding the presence of class 1 integrons in *E. fergusonii* except for a single study reported earlier^15^. The presence of different MGEs in association with ARGs, and VGs enabled the OTSVEF–60 strain to becoming a potential threat for public and animal health concern. Besides, *E. fergusonii* strains are frequently reported to become antibiotic resistant through the horizontal transfer of MGEs carrying the resistant traits^11^. Furthermore, the presence of insertion sequences (IS), and transposons (Tn) in the genome of OTSVEF–60 may also facilitate the dissemination of the resistance genes within the same or different species of bacteria^51^. The WGS of the OTSVEF–60 reveals CRISPR spacers present in the isolate might affect bacterial evolution, and interfere with the uptake of bacteriophage DNA carrying virulence genes, including toxin and ARGs as also predicted previously^52^. In addition, the genome of the OTSVEF–60 strain also possessed six prophages signatures in the chromosome, although it is not known if these phage immunities have some effect on such high level of metal/antibiotic resistance or vice-versa. However, it shows a possibility that this might facilitate the bacteria to overcome phage attacks.

In the KEGG analysis, a number of cellular processes like survival and adaptation promoting pathways have been observed within the genome. It can be inferred from its genetic potential of quite developed carbon and energy metabolisms that these systems might facilitate the bacterium to utilize diverse carbon source for energy production, and surviving diverse contaminated environment. Several KO’s linked to carbohydrate metabolism pathways or others were also present which are associated with an increase in fermentation products, and thus conferring stress tolerance in different environmental conditions leading to the fortification of resistant character of the strain^53^.

Our current study has shown that *E. fergusonii* can be a reservoir for the dissemination of ARGS, and a threat to the effective treatment of bacterial infections they cause. The presence of MGEs in the genome may further complicates the treatment of bacterial infections in clinical settings. This compete genome sequencing of the OTSVEF–60 isolate, and annotation of the WGS in search for detection of VGs, ARGs and associated metabolic functional genes added valuable information to the genetic features, and pathogenesis of the POEF strain to develop the prevention strategy and control measures. The results of this study open the door for further investigation of the associated mechanisms of virulence or pathogenicity, disease complication and/or treatment failure, and subsequent spreading of VGs and ARGs across the poultry settings of Bangladesh.

## Authors’ contributions

O.S. carried out the studies (sequencing, molecular and data analysis). O.S. and N.N.R. participated in drafting the manuscript. M. N.H edited and revised the final manuscript, prepared the final figures and tables. M.S. and M.A.H. developed the hypo thesis, supervised the whole work and helped to prepare and revise the manuscript. All authors read and approved the final manuscript.

## Acknowledgments

The authors would like to acknowledge Bangladesh Academy of Science – United States Department of Agriculture (BAS – USDA) for supporting this project. We would also like to acknowledge Bangabandhu Science & Technology Fellowship Trust for supporting Otun Saha as PhD student. We would like to further acknowledge University Grants Commission (UGC), Ministry of Science and Technology, Bangladesh for supporting reagents and equipment.

## Ethical approval

Ethical approval was granted from the Ethics Committee of the Faculty of Biological Sciences, University of Dhaka, Bangladesh who has approved the procedure under the Reference 71/Biol.Scs./2018-2019.

## References

1. Farmer JJ, Fanning GR, Davis BR, O’Hara CM, Riddle C, Hick-Brenner FW. et al. *Escherichia fergusonii* and *Enterobacter taylorae*, two new species of *Enterobacteriaceae* isolated from clinical specimens. J. Clin. Microbiol. 1985; 21, 77–81.

2. Adesina T, Nwinyi O, De N, Akinnola O, Omonigbehin E. First Detection of Carbapenem-Resistant Escherichia fergusonii Strains Harbouring Beta-Lactamase Genes from Clinical Samples. Pathogens 2019; 8(4), 164.

3. Glover B, Wentzel J, Jenkins A, Van Vuuren M. The first report of Escherichia fergusonii isolated from non-human primates, in Africa. One Health 2017; 4(3), 70–75. doi: 10.1016/j.onehlt.2017.05.001. PMID: 28616507; PMCID: PMC5454151.

4. Oh JY. et al. Isolation and epidemiological characterization of heat-labile enterotoxin-producing *Escherichia fergusonii* from healthy chickens. Vet. Microbiol. 2012; doi:10.1016/j.vetmic.2012.05.020.

5. Weiss ATA, Lübke-Becker A, Krenz M, van der Grinten. *E. Enteritis* and Septicemia in a Horse Associated With Infection by *Escherichia fergusonii*. J. Equine Vet. Sci. 2011; doi:10.1016/j.jevs.2011.01.005.

6. Hariharan H, Lopez A, Conboy G, Coles M, Muirhead T. Isolation of *Escherichia fergusonii* from the feces and internal organs of a goat with diarrhea. Canadian Vet. J. 2007; 48, 630–631.

7. Herráez P, Rodriguez AF, Espinosa A et al. Fibrinonecrotictyphlitis caused by *Escherichia fergusonii* in ostriches (Struthiocamelus). Avian Dis. 2005; 49, 167–169.

8. Touchon M. et al. Organised genome dynamics in the *Escherichia coli* species results in highly diverse adaptive paths. PLoS Genet. 2009; doi:10.1371/journal.pgen.1000344.

9. Fegan N, Barlow RS, Gobius KS. *Escherichia coli* O157 somatic antigen is present in an isolate of *E. fergusonii*. Curr. Microbiol. 2006; doi:10.1007/s00284-005-0447-6.

10. Lagacé-Wiens PR, Baudry PJ, Pang P, Hammond G. First Description of an Extended-Spectrum-ß-Lactamase-Producing Multidrug-Resistant *Escherichia fergusonii* Strain in a Patient with Cystitis. J. Clinical Microbiol. 2010; 48(6), 2301–2.

11. Forgetta V. et al. Pathogenic and multidrug-resistant *Escherichia fergusonii* from broiler chicken. Poult. Sci. 2012; doi:10.3382/ps.2011-01738.

12. Gaastra W, Kusters JG, van Duijkeren E, Lipman LJA. Escherichia fergusonii. Vet. Microbiol. doi:10.1016/j.vetmic.2014.04.016.

13. Fricke WF, McDermott PF, Mammel MK, Zhao SH, Johnson TJ, et al. Antimicrobial Resistance-Conferring Plasmids with similarity to virulence plasmids from avian pathogenic *Escherichia coli* strains in *Salmonella enterica* serovar Kentucky isolates from poultry. Appl. Environ. Microbiol. 2009; 75, 5963–5971.

14. Fernandez-Alarcon C, Singer RS, Johnson TJ. Comparative genomics of multidrug resistance-encoding IncA/C plasmids from commensal and pathogenic *Escherichia coli* from multiple animal sources. PLoS ONE 2011; 6, 23415. doi: 10.1371/journal.pone.0023415.

15. Simmons K, Islam MR, Rempel H, Block G, Topp E, Diarra MS. Antimicrobial resistance of *Escherichia fergusonii* isolated from broiler chickens. J. Food Protect. 79(6), 929–938.

16. Galetti R, Filho RACP, Ferreira JC, Varani AM, Darini ALC. Antibiotic resistance and heavy metal tolerance plasmids: The antimicrobial bulletproof properties of *Escherichia fergusonii* isolated from poultry. Infect Drug Resist. 2019; doi:10.2147/IDR.S196411.

17. Wragg P, La Ragione RM, Best A, Reichel R, Anjum MF, Mafura M, Woodward MJ. Characterisation of *Escherichia fergusonii* isolates from farm animals using an *Escherichia coli* virulence gene array and tissue culture adherence assays. Res. Vet. Sci. 2009; 86(1), 27–35.

18. Furtula V, Farrell EG, Diarrassouba F, Rempel H, Pritchard J, Diarra MS. Veterinary pharmaceuticals and antibiotic resistance of *Escherichia coli* isolates in poultry litter from commercial farms and controlled feeding trials. Poult. Sci. 2010; 89, 180–188.

19. Yu Z, Gunn L, Wall P, Fanning S. Antimicrobial resistance and its association with tolerance to heavy metals in agriculture production. Food Microbiol. 2017; doi:10.1016/j.fm.2016.12.009.

20. Momtaz S, Hossain MA. Occurrence of Pathogenic and Multidrug Resistant *Salmonella* spp. Biores. Comm. 2018; 4(2), 506–515.

21. Hoque MN, Istiaq A, Clement RA. et al. Insights into the Resistome of Bovine Clinical Mastitis Microbiome, a Key Factor in Disease Complication. Front. Microbiol. 2020; 11, 860

22. Saha O, Hoque MN, Islam OK. et al. Multidrug-Resistant Avian Pathogenic *Escherichia coli* Strains and Association of Their Virulence Genes in Bangladesh. bioRxiv. 2020; doi: https://doi.org/10.1101/2020.06.30.180257.

23. Slatko BE, Gardner AF, Ausubel FM. Overview of next□generation sequencing technologies. Current Prot. Mol. Biol. 2018; 122(1), e59.

24. Bolger AM, Lohse M, Usadel B. Trimmomatic: a flexible trimmer for Illumina sequence data. Bioinformatics 2014; 30(15), 2114–2120.

25. Bankevich A, Nurk S, Antipov D, Gurevich AA. et al. SPAdes: a new genome assembly algorithm and its applications to single-cell sequencing. J. Comput. Biolo. 2012; 19(5), 455–477. Doi:10.1089/cmb.2012.0021.

26. Darling AC, Mau B, Blattner FR, Perna NT. Mauve: multiple alignment of conserved genomic sequence with rearrangements. Genome Res. 2004; 14(7), 1394–1403 DOI 10.1101/gr.2289704.

27. Aziz RK, Bartels D, Best AA, DeJongh M, Disz T, Edwards RA. et al. The RAST Server: rapid annotations using subsystems technology. BMC Genomics 2008; 9, 75. doi: 10.1186/1471-2164-9-75.

28. Moriya Y, Itoh M, Okuda S, Yoshizawa AC, Kanehisa M. KAAS: an automatic genome annotation and pathway reconstruction server. Nucleic Acids Res. 2007; 35, W182–W185 DOI 10.1093/nar/gkm321.

29. Alikhan NF, Petty NK, Zakour NLB, Beatson SA. BLAST Ring Image Generator (BRIG): simple prokaryote genome comparisons. BMC Genomics 2011; 12(1), 402.

30. Goris J, Konstantinidis KT, Klappenbach JA, Coenye T, Vandamme P, Tiedje, J M. DNA-DNA hybridization values and their relationship to whole-genome sequence similarities. Int. J. Syst. Evol. Micr. 2007; 57, 81–91.

31. Lowe TM, Eddy SR. TRNAscan-SE: a program for improved detection of transfer RNA Genes in genomic sequence. Nucleic Acids Res. 1997; 25(5), 955–964.

32. Lagesen K, Hallin P, Rødland EA, Stærfeldt HH, Rognes T, Ussery DW. RNAmmer: consistent and rapid annotation of ribosomal RNA genes. Nucleic Acids Res. 2017; 35(9), 3100–3108.

33. Joensen KG, Scheutz F, Lund O, Hasman H, Kaas RS, Nielsen EM. et al. Real-time whole-genome sequencing for routine typing, surveillance, and outbreak detection of verotoxigenic *Escherichia coli*. J. Clin. Microbiol. 2014; 52, 1501–1510.

34. Zhang Q, Ye Y. Not all predicted CRISPR–Cas systems are equal: isolated cas genes and classes of CRISPR like elements. BMC Bioinformatics 2017; 18, 92. doi: 10.1186/s12859-017-1512.

35. Siguier P, Pérochon J, Lestrade L, Mahillon J, Chandler M. ISfinder: the reference centre for bacterial insertion sequences. Nucleic Acids Res. 2006; 34, D32–D36.

36. Medema MH, Blin K, Cimermancic P, De Jager V, Zakrzewski P, Fischbach MA, Weber T, Takano E, Breitling R. antiSMASH: rapid identification, annotation and analysis of secondary metabolite biosynthesis gene clusters in bacterial and fungal genome sequences. Nucleic Acids Res. 2011;39, W339–W346 DOI 10.1093/nar/gkr466.

37. Kumar S, Stecher G, Tamura K. MEGA7: Molecular evolutionary genetics analysis version 7.0 for bigger datasets. Mol. Biol. Evol. 2016; 33, 1870–1874.

38. Letunic I, Bork P. Interactive tree of life (iTOL) v3: an online tool for the display and annotation of phylogenetic and other trees. Nucleic Acids Res. 2016; 44(1), 242–5.

39. Adékambi T, Colson P, Drancourt M. rpoB-Based Identification of Nonpigmented and Late-Pigmenting Rapidly Growing Mycobacteria. J. Clin. Microbiol. 2003; doi:10.1128/JCM.41.12.5699-5708.2003.

40. Puttamreddy S, Cornick NA, Minion FC. Genome-wide transposon mutagenesis reveals a role for pO157 genes in biofilm development in *Escherichia coli* O157: H7 EDL933. Infect. Immun. 2011; 78, 2377–2384.

41. Karczmarczyk M, Abbott Y. Walsh C, Leonard N, Fanning S. Characterization of multidrug-resistant *Escherichia coli* isolates from animals presenting at a university veterinary hospital. Appl. Environ. Microbiol. 2011; 77, 7104–7112

42. White PA, Rawlinson WD. Current status of the *aadA* and *dfr* gene cassette families. J. Antimicrob. Chemother. 2001;47, 495–496.

43. Nagakubo S, Nishino K, Hirata T. The Putative Response Regulator BaeR Stimulates Multidrug Resistance of *Escherichia coli* via a Novel Multidrug Exporter System, MdtABC The Putative Response Regulator BaeR Stimulates Multidrug Resistance of Escherichia coli via a Novel Multidrug Exporter. J. Bacteriol. 2002;184(15), 4161–4167.

44. Yung PY. et al. Global transcriptomic responses of *Escherichia coli* K-12 to volatile organic compounds. Sci. Rep. 2016; 6, 19899.

45. Bohnert JA, Schuster S, Fähnrich E, Trittler R, Kern WV. Altered spectrum of multidrug resistance associated with a single point mutation in the *Escherichia coli* RND-type MDR efflux pump YhiV (MdtF). J. Antimicrob. Chemother. 2007;59, 1216–1222.

46. Lemire JA, Harrison JJ, Turner RJ. Antimicrobial activity of metals: mechanisms, molecular targets and applications. Nat. Rev. Microbiol. 2013; 11(6), 371–384.

47. McIntosh D, Cunningham M, Ji B, Fekete FA. et al. Transferable, multiple antibiotic and mercury resistance in Atlantic Canadian isolates of *Aeromonas salmonicida* subsp. salmonicida is associated with carriage of an IncA/C plasmid similar to the *Salmonella* enterica plasmid pSN254. J. Antimicrob. Chemoth. 2008; 61(6), 1221–1228.

48. La Mendola D, Giacomelli C, Rizzarelli E. Intracellular Bioinorganic Chemistry and Cross Talk Among Different-Omics. Curr. Top Med. Chem. 2016; 16, 3103–3130.

49. Hellweger FL. *Escherichia coli* adapts to tetracycline resistance plasmid (pBR322) by mutating endogenous potassium transport: in silico hypothesis testing. FEMS Microbiol. Ecol. 2013; 83, 622–631.

50. Rowe-Magnus DA, Mazel D. The role of integrons in antibiotic resistance gene capture. Int. J. Medical Microbiol. 2002; 292(2), 115–125.

51. Partridge SR, Kwong SM, Firth N, Jensen SO. Mobile genetic elements associated with antimicrobial resistance. Clin. Microbiol. Rev. 2018; 31(4), 88–17.

52. Louwen R, Staals RH, Endtz HP, van Baarlen P, van der Oost J. The role of CRISPR-Cas systems in virulence of pathogenic bacteria. Microbiol. Mol. Biol Rev. 2014; 78(1), 74–88.

53. Hoque MN, Istiaq A, Clement RA. et al. Metagenomic deep sequencing reveals association of microbiome signature with functional biases in bovine mastitis. Sci. Rep. 2019; 9, 13536.

