## Supplementary Figure Legends for "Genome-wide Genetic Marker Analysis and Genotyping of *Escherichia fergusonii* strain OTSVEF–60"

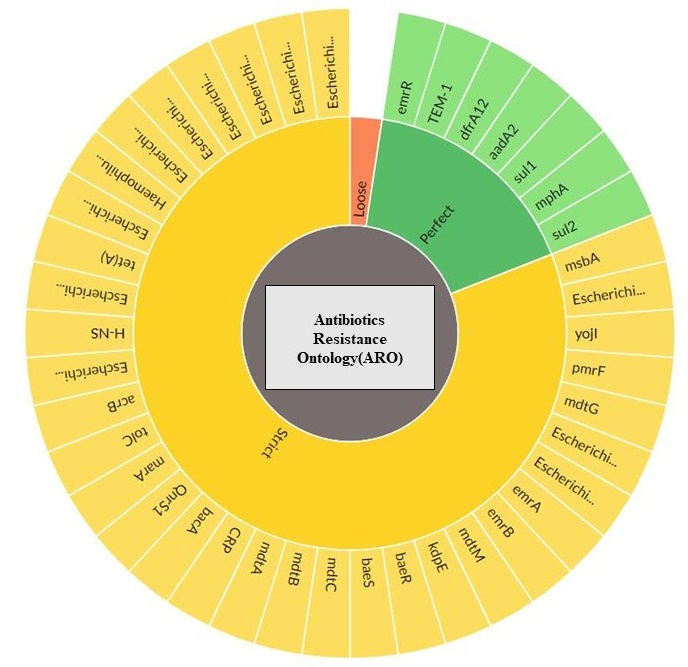

Supplementary Fig.1. Antibiotic Resistance Ontology (ARO) of POEF OTSVEF –60based on RGI (resistance gene identifier) criteria (perfect, strict, complete genes only) using RGI 4.2.0 (https://card.mcmaster.ca/analyze/rgi).

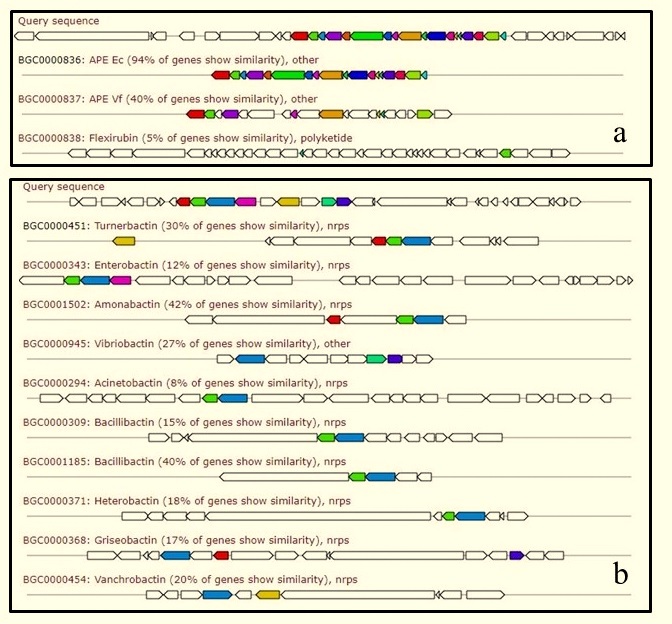

Supplementary Fig. 2. Metabolite Analysis of POEF OTSVEF-60 by Anti-SMASH. a) Arylpolyene Gene Clusters Comprise the Largest Known biosynthetic [gene clusters](https://www.sciencedirect.com/topics/biochemistry-genetics-and-molecular-biology/gene-cluster) (BGCs) Family; b) non-ribosomal peptide synthase (NRPS) Gene Clusters Comprise the Largest Known biosynthetic [gene clusters](https://www.sciencedirect.com/topics/biochemistry-genetics-and-molecular-biology/gene-cluster) (BGCs).
