## Supplementary Tables for "Genome-wide Genetic Marker Analysis and Genotyping of *Escherichia fergusonii* strain OTSVEF–60"

**Supplementary Table 1:** Sequence types (ST) of *E. fergusonii* OTSVEF –60 strains isolated from poultry feces determined using MLST 2.0 (Multi-Locus Sequence Typing) and verified by (http://enterobase.warwick.ac.uk.).

| **Locus** | **Identity** | **Coverage** | **Alignment Length** | **Allele Length** | **Gaps** | **Allele** |
| --- | --- | --- | --- | --- | --- | --- |
| *adk* | 100 | 100 | 536 | 536 | 0 | *adk*_491 |
| *fumC* | 99.7868 | 100 | 469 | 469 | 0 | *fumC*_204* |
| *gyrB* | 100 | 100 | 460 | 460 | 0 | *gyrB*_701 |
| *icd* | 99.6139 | 100 | 518 | 518 | 0 | *icd*_319* |
| *mdh* | 100 | 100 | 452 | 452 | 0 | *mdh*_401 |
| *purA* | 100 | 100 | 478 | 478 | 0 | *purA*_40 |
| *recA* | 100 | 100 | 510 | 510 | 0 | *recA*_391 |

Notes: * alleles with less than 100% identity found

**Supplementary Table 2:** Characteristic features of CRISPR-Cas system in POEF OTSVEF-60 strains isolated from poultry feces.

| **Locus** | **Sub-type** | **Cas Proteins** | **No. of Repeats** | **Average length of Repeats** | **No of spacer** | **Average length of Spacer** | **Suspicious *cas* genes and/or false-CRISPRs** |
| --- | --- | --- | --- | --- | --- | --- | --- |
| 1 | Type-1,1-A | Csa3,DEDDH,Cas3 | ---- | ---- | ---- | ---- | + |
|  |  | ---- | 3 | 22 | 2 | 38 |  |
| 2 | Type-1,1-A | CasR,Csa3 | ----- | ---- | --- | --- |  |
| 3 | Type-1,1-E | Cas1,Cas2,Cas6e,Cas7,Cas5,Cas8e,Cas3,Csa2gr11 | 42 | 29 | 40 | 32 |  |

**Supplementary Table 3**: Secondary Metabolite Analysis of POEF OTSVEF-60 by Anti-SMASH

| **Input Accession Number** | **Position** | **Gene Cluster Type** | **Detected gene Cluster genes In the Database** |
| --- | --- | --- | --- |
| c00003_NODE_3_.. | NODE_3_length_203303_cov_90.237010 | arylpolyene | ctg3_(136-172) |
| c00039_NODE_39.. | NODE_39_length_28312_cov_75.502679 | NRPS | ctg39_(1-33) |

**Supplementary Table 4**: IS elements in the POEF OTSVEF-60 genome.

| **IS element** | **IS Family** | **Sources** | **Product** | **Length(bp)** | **Integrity** |
| --- | --- | --- | --- | --- | --- |
| Tn2 | Tn3 | *Escherichia coli* | Transposase | 4950 | complete |
| Tn3 | Tn3 | *Salmonella enterica* | Transposase | 4948 | complete |
| ISKpn19 | ISKra4 | *Klebsiella pneumoniae* | Hypothetical protein | 2851 | truncated |
| TnAs3 | Tn3 | *Aeromonassalmonicida* | transposase | 18735 | truncated |
| TnAs1 | Tn3 | *Aeromonassalmonicida* | Hypothetical protein | 6694 | truncated |
| IS609 | IS200/IS605 | *Escherichia coli* | Transposase | 1748 | complete |
| ISEc81 | IS110 | *Escherichia coli* | Transposase | 1385 | complete |
| IS2 | IS3 | *Escherichia coli* | Transposase | 1331 | truncated |
| ISEc17 | IS3 | *Escherichia coli* | Transposase | 1258 | truncated |
| IS3 | IS3 | *Escherichia coli* | Transposase | 1258 | truncated |
| ISKpn26 | IS3 | *Klebsiella pneumoniae* | Transposase | 1258 | truncated |
| ISEc51 | ISKra4 | *Escherichia sp.* | Transposase | 2857 | truncated |
| IS3F | IS3 | *Escherichia fergusonii* | Transposase | 1258 | truncated |
| IS5075 | IS110 | *Escherichia coli* | Transposase | 1327 | complete |
| ISEc1 | ISAs1 | *Escherichia coli* | Transposase | 1291 | truncated |
| ISEc5 | ISAs1 | *Escherichia coli* | Transposase | 1291 | truncated |
| ISPa40 | Tn3 | *Pseudomonas aeruginosa* | Chromate resistance protein; Transposase | 6592 | truncated |
| ISEc42 | IS200/IS605 | *Escherichia coli* | Accessory Gene | 1291 | complete |
| IS3H | IS3 | *Shigelladysenteriae* | Transposase | 1261 | truncated |
| IS6100 | IS6 | *Mycobacterium fortuitum* | Transposase | 880 | complete |
| IS26 | IS6 | *Proteus vulgaris* | Transposase | 820 | complete |
| IS15DII | IS6 | *Salmonella panama* | Transposase | 820 | complete |
| IS15DIV | IS6 | *Salmonella typhimurium* | Transposase | 820 | complete |
| IS15 | IS6 | *Salmonella panama* | Transposase | 1648 | complete |
| IS15DI | IS6 | *Salmonella panama* | Transposase | 820 | complete |
| IS5 | IS5 | *Escherichia coli* | Transposase | 1195 | truncated |
| IS5D | IS5 | *Escherichia coli* | Transposase | 1283 | truncated |
| IS4321 | IS110 | *----* | Transposase | 1327 | truncated |
| IS4321L | IS110 | *Enterobacter aerogenes* | Transposase | 1326 | complete |
| IS4321R | IS110 | *Enterobacter aerogenes* | Transposase | 1326 | complete |
| IS5B | IS5 | *Enterobacter aerogenes* | Transposase | 1207 | truncated |
| IS4 | IS4 | *Escherichia coli* | Transposase | 1426 | truncated |
| ISEc27 | IS3 | *Escherichia coli* | Transposase | 1321 | truncated |
| IS1G | IS1 | *Escherichia coli* | Transposase | 768 | truncated |
| IS1A | IS1 | *Escherichia coli* | Transposase | 768 | truncated |
| IS1R | IS1 | *Escherichia coli* | Transposase | ---- | truncated |
| IS1SD | IS1 | *Shigelladysenteriae* | Transposase | 768 | truncated |
| IS1B | IS1 | *Escherichia coli* | Transposase | 768 | truncated |
| IS1S | IS1 | *Shigellasonnei* | Transposase | 768 | truncated |
| ISPa38 | Tn3 | *Pseudomonas aeruginosa* | Hypothetical protein;Tn3 resolvase | 6455 | complete |
| ISShes11 | Tn3 | *Shewanella sp.* | Transposase | 7669 | truncated |
| ISEc44 | IS200/IS605 | ***----*** | Transposase | 1879 | complete |
